# Chimeric Genes Revealed in the Polyploidy Fish Hybrids of *Carassius cuvieri* (Female) × *Megalobrama amblycephala* (Male)

**DOI:** 10.1101/082222

**Authors:** Fangzhou Hu, Chang Wu, Yunfan Zhou, Shi Wang, Jun Xiao, Yanhong Wu, Kang Xu, Li Ren, Qingfeng Liu, Wuhui Li, Ming Wen, Min Tao, Qinbo Qin, Rurong Zhao, Kaikun Luo, Shaojun Liu

## Abstract

The genomes of newly formed natural or artificial polyploids may experience rapid gene loss and genome restructuring. In this study, we obtained tetraploid hybrids (4n=148, 4nJB) and triploid hybrids (3n=124, 3nJB) derived from the hybridization of two different subfamily species *Carassius cuvieri* (♀, 2n = 100, JCC) and *Megalobrama amblycephala* (♂, 2n = 48, BSB). Some significant morphological and physiological differences were detected in the polyploidy hybrids compared with their parents. To reveal the molecular traits of the polyploids, we compared the liver transcriptomes of 4nJB, 3nJB and their parents. The results indicated high proportion chimeric genes (31 > %) and mutated orthologous genes (17 > %) both in 4nJB and 3nJB. We classified 10 gene patterns within three categories in 4nJB and 3nJB orthologous gene, and characterized 30 randomly chosen genes using genomic DNA to confirm the chimera or mutant. Moreover, we mapped chimeric genes involved pathways and discussed that the phenotypic novelty of the hybrids may relate to some chimeric genes. For example, we found there is an intragenic insertion in the K+ channel *kcnk5b*, which may be related to the novel presence of the barbels in 4nJB. Our results indicated that the genomes of newly formed polyploids experienced rapid restructuring post-polyploidization, which may results in the phenotypic and phenotypic changes among the polyploidy hybrid offspring. The formation of the 4nJB and 3nJB provided new insights into the genotypic and phenotypic diversity of hybrid fish resulting from distant hybridization between subfamilies.

## Introduction

Distant hybridization is defined as above-specific or interspecific crossing, and is a useful strategy to produce hybrid offspring with altered genotypes and phenotypes [1], or with different ploidies [2–4]. Allopolyploids, resulting from interspecific which combinations of two or more differentiated genomes, are more prevalent in plants than in vertebrates [5–7]. Previous research has suggested that the reasons for this difference include genome shock or dramatic genomic restructuring[6]; environmental fluctuations[7]; barriers to sex determination, physiological and developmental constraints; especially nuclear–cytoplasmic interactions and related factors[8]. However, to date, the mechanisms underlying this difference between plants and vertebrates are unknown.

In the wild, research on polyploidy is limited by the the availability of plant and animal materials, because allopolyploids were usually formed hundreds or even thousands of years ago, and their original diploid parental species are often extinct or unknown [9, 10]. Thus, as model systems, synthetic allopolyploids represent excellent genetic materials to study and characterize drastic genomic changes at early evolutionary stages. Fish are one of the minority of vertebrates in which polyploidy clearly exists (amphibians and reptiles also show polyploidy)[11]. Fish chromosomes display plasticity, and thus it is easier to produce synthetic allopolyploids using distant hybridization between fishes.

In the fish catalog, the Japanese crucian carp (*Carassius cuvieri*) (JCC) with 100 chromosomes belongs to the Cyprininae subfamily, and the blunt snout bream (*Megalobrama amblycephala*) (BSB) with 48 chromosomes belongs to the Cultrinae subfamily[12]. In this study, we successfully obtained tetraploid (4n=148, 4nJB) and triploid (3n=124, 3nJB) hybrids by crossing JCC ♀ and BSB ♂. Interestingly, both 4nJB and 3nJB showed phenotypic differences compared with their parents. Such as 4nJB hybrids have one pair barbels but their parents have no barbels, and 3nJB have slightly prominent eyes compared with their parents. In addition, the 4nJB are bisexual fertility, but 3nJB are sterile. Thus, these hybrids are an appropriate model to investigate the relationships between phenotypes and genotypes in hybrid fish, and to study instantaneous allopolyploidization and the crucial changes that immediately follow hybridization.

Previous studies indicated that hybridization followed by allopolyploidization cause drastic genetic and genomic imbalances, including chromosomal rearrangements[13, 14]; transpositions[15]; deletions and insertions[16]; dosage imbalances[17]; a high rate of DNA mutations and recombinations[18, 19]; and other non-Mendelian phenomena[20, 21]. However, little is known about the molecular mechanisms underlying these changes in new allopolyploids. The next-generation sequencing data provided in this study provides resources to address such questions as how hybridization affects gene variation that could lead to biological characters in nascent hybrid fish. In this study, we performed RNA-seq on liver tissue samples from 4nJB, 3nJB and their parents (JCC and BSB) to reveal the molecular traits of the polyploids hybrid fish. This is the first report of tetraploid and triploid hybrids produced by crossing female Japanese crucian carp and male blunt snout bream. The production of two new hybrid offspring has significance in fish genetic breeding and evolutionary studies.

## Materials and Methods

### Ethics statement

Administration of Affairs Concerning Animal Experimentation Guidelines states that approval from the Science and Technology Bureau of China and the Department of Wildlife Administration is not necessary when the fish in question are not rare or not near extinction (first-class or second-class state protection level). Therefore, approval is not required for the experiments conducted in this study.

### Formation of triploid and tetraploid hybrids

Individuals of Japanese crucian carp (JCC), blunt snout bream (BSB), the tetraploid F_1_ hybrids (4nJB) and triploid F_1_ hybrids (3nJB) of female JCC × male BSB were obtained from the Engineering Research Center of Polyploid Fish Breeding and Reproduction of the State Education Ministry, China, located at Hunan Normal University. During the reproductive seasons (from April to June each year) in 2013, 2014 and 2015, 20 mature females and 20 mature males of both JCC and BSB were chosen as parents. The crossings were performed in four groups. In the first group, the JCC was used as the female parent, and the BSB was used as the male parent. In the second group, the female and male parents were reversed. In the third group, the male JCC and female JCC were allowed to self-mate (crossing of brothers and sisters). In the fourth group, male and female BSB were used for self-mating. The mature eggs and sperm of JCC and BSB were fertilized and the embryos were developed in culture dishes at a water temperature of 19–20°C. In each group, 5000 embryos were taken at random for the examination of the fertilization rate (number of embryos at the stage of gastrula/number of eggs), the hatching rate (number of hatched fry/number of eggs) and early survival rate (number of surviving fry/ number of hatched fry). The hatched fry were transferred to the pond for further culture.

### Measurement morphological traits of triploid and tetraploid hybrids

The ploidy levels of F_1_ hybrids were confirmed by measuring DNA content, forming chromosomal karyotypes and using the erythrocyte nuclear volume. The methods refer to previous study[22, 23]. The morphology and fertility of 4nJB and 3nJB were also examined. The examined measurable traits included the average values of the whole length (WL), the body length (BL) and width BW, the head length (HL) and width (HW), and the tail length (TL) and width (TW). Using these measurements, the following ratios were calculated: WL/B, BL/BW, BL/HL, HL/HW, TL/TW and BW/HW. The examined countable traits included the number of dorsal fins, abdominal fins, anal fins, lateral scales, and upper and lower lateral scales. For both measurable and countable data, we used SPSS software to analyze the covariance of the data between the hybrid offspring and their parents. At 15 months old, 20 4nJB and 20 3nJB individuals were randomly sampled to examine their gonad development by histological sectioning. The methods refer to previous study[24]. The gonadal stages were classified according to Liu’s standard series for cyprinid fish[25].

### Transcriptome sequencing and analysis

#### cDNA library construction, transcriptome sequencing and quality control

At 15 months old, three fish of each type were picked randomly and euthanized using 2-phenoxyethanol (Sigma, USA) before being dissected. Liver tissues were excised and immediately placed in RNALater (Ambion, USA), following the manufacturer’s instructions, for storage. RNA was isolated according to the standard Trizol protocol (Qiagen, Valencia, CA, USA), and quantified using an Agilent 2100 Bioanalyzer (Agilent, Santa Clara, CA, USA). The RNA used for subsequent cDNA library construction was a pooled sample from the liver tissue of the three different individuals.

The four cDNA libraries representing each type of fish (4nJB, 3nJB, JCC and BSB) were constructed from 2 μg of mRNA. Each library was sequenced using an Illumina HiSeq™ 2000/2500. From the generated raw reads, the read adaptors and low-quality reads were removed and the clean reads from each library were examined using software FastQC. Transcriptome de novo assembly was carried out using a short-reads assembly program (Trinity) [26], with three independent software modules called Inchworm, Chrysalis, and Butterfly. Principal component analysis (PCA) of 4 liver transcriptomes was applied to examine the contribution of each transcript to the separation of the classes[27, 28].

Contig annotation was performed using five public databases (nonredundant (Nr); Swiss-Prot; Kyoto Encyclopedia of Genes and Genomes (KEGG); Clusters of Orthologous Groups (COG) and Gene Ontology (GO)). BLASTX alignment (e-value ≤ 1e^−6^) between contigs and protein databases was performed, and the best-aligned results were used to decide the sequence direction of the contigs. After screening the sequences (alignment length ≤ 100 bp), accession numbers of the genes were obtained from the BLASTX results. Then, Go terms of annotated sequences were obtained using Ensembl BioMart[29]. WEGO software was used to analyze the GO annotation [30]. To identify putative orthologs between the hybrid offspring and the parents, the sequences from the four assembled transcriptomes were aligned using reciprocal BLAST (BLASTN) hit with an e-value of 1e^−20^ [31]. Four sequences were defined as orthologs if each of them was the best hit of the other and if the sequences were aligned over 300 bp. Meanwhile, the nucleotide sequences were aligned using the BioEdit program (version 7.0.9) [32].

### Variation and detection of chimeric patterns

High-quality reads were remapped to BSB reference genome with Burrows-Wheeler Alignment tool (BWA) [33] to detect variants among 4nJB, 3nJB, JCC and BSB, and their distributions. Because divergence within most shared copies of both JCC and BSB was less than 5%, the maximum mismatch of hits was set as 5 per 100 bp read. This parameter setting (maximum mismatch of 5%) ensured that reads from half of the orthologs were mapped to the genome as the mapping ratio of the blunt snout bream is about 50%. After obtaining the BAM files, we recorded the mapped region of each sample on the reference genome. Variations from regions overlapped by both parents and tetraploid (triploid) hybrids were extracted from the alignments using both mpileup from the SAMtools package[34], and the GATK [35–37] pipeline for RNA-seq. Candidate variations were filtered based on a variation-quality score ≥20, and depth >3 reads. VCFtools [38] was used to compare variations among both parents and tetraploid (triploid) hybrids identified by both methods.

The distribution patterns of variations in tetraploid and triploid hybrids compared to both parents were analyzed and the distributions of chimeric loci were retained for downstream analysis. Mutation patterns were defined as follows: first, parents-variants, where the sequence was same as one parent but different from the other parent; second, offspring-specific-variants, such as DNA insertions, DNA deletions or locus mutations; third, chimeric-variants, single or multiple fragments consisting of continuous, alternating variations from parents-specific variants. Within a gene-region, several segmental fragments potentially consisted of different exons. Thus, offspring mutations were associated with segments. Genes with single or multiple exons derived only from one parent were classified as being of maternal-origin or paternal-origin (patterns 1 and 2). Genes of offspring with specific variations but no chimeric patterns were identified as having mutations (patterns 3–8). If parent-specific variations aligned alternately within a continuous fragment, or occurred alternately in non-continuous fragments, the genes was classified as a chimera containing a parental crossing hotspot (patterns 9–10). Only genes with fragments per kilobase of transcript per million mapped reads (FPKM) values > 0.3 were classified by pattern. Redundant genes (the same gene name and pattern of variation) in the Japanese crucian carp-based analyses were removed.

### PCR validation for 30 chimeric genes and mutated *kcnk5b* gene

Artificial chimeras might have existed within some 4nAT and 3nAT sequences. To verify the accuracy of sequencing assembly and estimate the genomic changes, thirty genes were chosen randomly from each pattern to validate the chimeric pattern. The genomic DNAs were extracted from the liver tissue of JCC, BSB and their hybrid offspring by routine approaches[39]. Polymerase chain reaction (PCR) primers were designed based on the conserved regions of homologous gene from JCC, BSB, 4nJB and 3nJB (Table S1). The PCR was performed in a volume of 50 μl with approximately 160 ng of genomic DNA, 3 mm of MgCl_2_, 400 mm of each dNTP, 0.6 mm of each primer, and 1.8 unit of Taq polymerase (Takara). The cycling program included 35 cycles of 94°C for 30 sec, 52–59°C for 30 sec, and 72°C for 1–5 min, with a final extension of 10 min at 72°C. Amplification products were separated on a 1.2% agarose gel using TBE buffer. PCR products within the expected size range were extracted and purified using a gel extraction kit (Sangon, Shanghai, China) and ligated into the pMD18-T vector. The plasmids were amplified in DH5α. The inserted DNA fragments subjected to Sanger sequencing using an ABI 3730 DNA Analyzer (Applied Biosystems, Carlsbad, CA, USA). To determine sequence homology and variation among the fragments, the sequences were aligned using BioEdit[32] and ClustalW2 (see URLs: www.ebi.ac.uk/Tools/es/cgi-bin/clustalw2). In addition, those validated genes identified from blunt snout bream reference-genome were combined to detect the structural changes.

## RESULTS

### Formation and measurement morphological traits of triploid and tetraploid hybrids

Each cross combination had a different of fertilization rate, hatching rate and early survival rate (Table 1). Allopolyploid hybrid fish were only generated from female *C. cuvieri* × male M. *amblycephala* (Fig. 1 C and D). The ploidy levels of F_1_ hybrids were confirmed by measuring their DNA content (Table 2), counting chromosomal numbers, chromosomal karyotype analysis (Table 3, Figure 1 E and F), and erythrocyte nuclear volume determination (Table 4, Figure 1 I and J). The results showed that two types of F_1_ hybrids were obtained: tetraploid (4n = 148) and triploid (3n = 124) hybrids.

**Table 1.** Fertilization rate, hatch rate and early survival rate of the different groups

| Fish type | Fertilization rate (%) | Hatch rate (%) | Early survival rate (%) |
| --- | --- | --- | --- |
| BSB × BSB | 99.5 | 89.6 | 83.1 |
| JCC × JCC | 98.4 | 86.2 | 82.5 |
| JCC × BSB | 86.1 | 66.4 | 75.7 |
| BSB × JCC | 85.7 | 0.5 | 0 |

**Table 2.** Mean DNA content of BSB, JCC, 3nJB and 4nJB hybrids

| Fish type | DNA<br>content | Ratio |  |
| --- | --- | --- | --- |
|  |  | Observed | Expected |
| BSB | 68.8 |  |  |
| JCC | 94.72 |  |  |
| 3nJB | 124.3 | $3nJB/(JCC+0.5BSB)=0.96^a$ | 1 |
| 4nJB | 157.43 | $4nJB/(JCC+BSB)=0.96^a$ | 1 |

<sup>a</sup> The observed ratio was not significantly different ( $P > 0.05$ ) from the expected ratio.

**Table 3.** Examination of chromosome number in BSB, JCC, 3nJB and 4nJB hybrids

| Fish type | No. in<br>metaphase | Distribution of chromosome number |  |  |  |  |  |  |  |
| --- | --- | --- | --- | --- | --- | --- | --- | --- | --- |
|  |  | <48 | 48 | <100 | 100 | <124 | 124 | <148 | 148 |
| BSB | 200 | 9 | 191 |  |  |  |  |  |  |
| JCC | 200 |  |  | 15 | 185 |  |  |  |  |
| 3nJB | 200 |  |  |  |  | 38 | 162 |  |  |
| 4nJB | 200 |  |  |  |  |  |  | 31 | 179 |

**Table 4.** Mean erythrocyte nuclear volume measurements for BSB, JCC, 3nJB and 4nJB hybrids

| Fish type | Major axis($\mu\text{m}$) | Minor axis($\mu\text{m}$) | Volume ($\mu\text{m}^3$) | Volume Ratio | |
| --- | --- | --- | --- | --- | --- |
|  |  |  |  | Observed | Expected |
| BSB | $5.07 \pm 0.47$ | $2.44 \pm 0.80$ | $15.76 \pm 1.92$ | | |
| JCC | $5.09 \pm 0.27$ | $2.92 \pm 0.20$ | $22.66 \pm 2.42$ | | |
| 3nJB | $6.86 \pm 0.70$ | $3.16 \pm 0.33$ | $35.76 \pm 1.67$ | $3\text{nJBL}/(\text{JCC}+0.5\text{BSB})=1.17^{\text{a}}$ | 1 |
| 4nJB | $7.38 \pm 0.56$ | $3.24 \pm 0.34$ | $40.53 \pm 1.42$ | $4\text{nJBL}/(\text{JCC}+\text{BSB})=1.05^{\text{a}}$ | 1 |

<sup>a</sup> The observed ratio was not significantly different ( $P > 0.05$ ) from the expected ratio.

**Figure 1.**
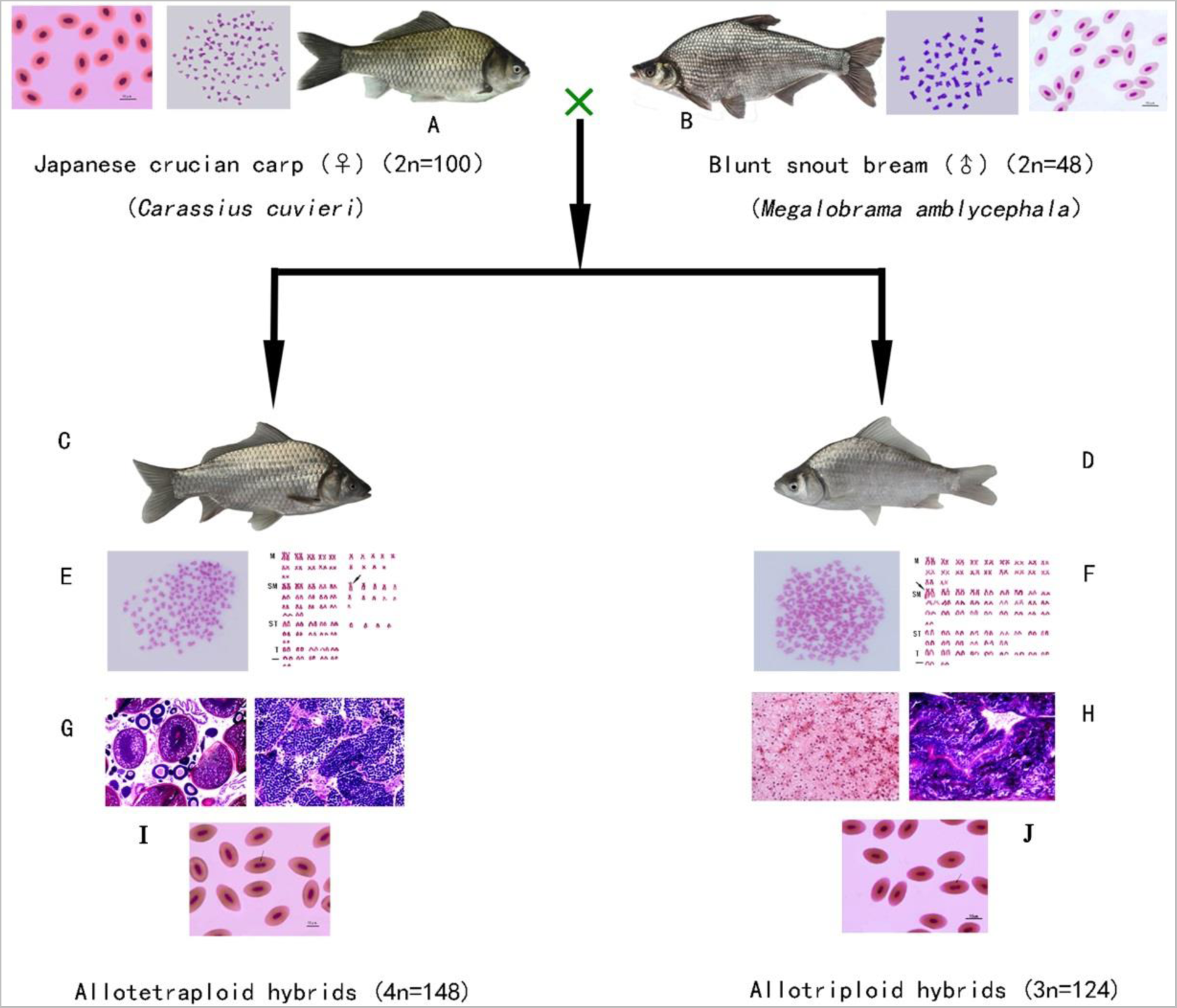
The chromosomal trait, gonadal development, erythrocytes and appearance of Japanese crucian carp (2n = 100), blunt snout bream (2n = 48), and their tetraploid and triploid F_1_ hybrid offspring. Appearance of JCC, BSB, 4nJB and 3nJB hybrids: (A) JCC; (B)BSB (C) 4nJB; (D) 3nJB. Chromosome and Karyotypes in 4nJB and 3nJB: (E) The metaphase chromosome spreads of 4nJB (4n =148); (F) The metaphase chromosome spreads of 3nJB (3n =124). The ovarian and testis microstructure of 3nJB and 4nJB: (G) 4nJB; (H) 3nJB. Erythrocytes of the 4nJB and 3nJB: (I) Normal erythrocytes with one nucleus and unusual erythrocytes with two nuclei (arrows) in a 4nJB; (J) Normal erythrocytes with one nucleus and unusual erythrocytes with two nuclei (arrows) in a 3nJB.

Measurable and countable traits were examined in each sample of BSB, JCC, 4nJB and 3nJB hybrids (Figure 1, Tables 5 and Tables 6). For morphological traits, the 4nJB (Figure 1C) and 3nJB (Figure 1D) showed obvious differences from JCC (Figure 1A) and BSB (Figure 1B). Most of the morphological data in 4nJB and 3nJB were significantly different from those in JCC and BSB (Tables 5 and Tables 6), suggesting that the traits varied in the allopolyploid hybrid offspring. Something interesting phenotypic variation occurred in hybrid offspring; for example, 4nJB have one pair barbels but their parents have no barbels; and the eyes of 3nJB are slightly prominent compared with their parents.

**Table 5.** Comparison of the measurable traits between the hybrid offspring and JCC and BSB

| Fish type | WL/BL | BL/BW | BL/HL | HL/HW | TL/TW | BW/HW |
| --- | --- | --- | --- | --- | --- | --- |
| BSB | $1.19 \pm 0.03$ | $2.37 \pm 0.03$ | $4.75 \pm 0.04$ | $1.14 \pm 0.03$ | $1.08 \pm 0.04$ | $2.09 \pm 0.04$ |
| JCC | $1.24 \pm 0.02$ | $2.22 \pm 0.15$ | $3.70 \pm 0.21$ | $1.17 \pm 0.06$ | $0.81 \pm 0.01$ | $1.78 \pm 0.09$ |
| 3nJB | $1.16 \pm 0.03$ | $2.28 \pm 0.13$ | $3.80 \pm 0.25$ | $1.03 \pm 0.07$ | $1.05 \pm 0.08$ | $1.72 \pm 0.11$ |
| 4nJB | $1.21 \pm 0.03$ | $2.35 \pm 0.12$ | $3.81 \pm 0.14$ | $1.12 \pm 0.06$ | $1.02 \pm 0.08$ | $1.83 \pm 0.10$ |

**Table 6.** Comparison of the countable traits between the hybrid offspring and JCC and BSB

| Fish type | No. of lateral scales | No. of upper lateral scales | No. of lower lateral scales | No. of dorsal fins | No. of abdominal fins | No. of anal fins |
| --- | --- | --- | --- | --- | --- | --- |
| BSB | 50.90 ± 0.91 (52–49) | 9.65 ± 0.49 (9–10) | 10.05 ± 0.69 (9–11) | III + 25.85 ± 0.59 (III 25–27) | 9.00 ± 0.55 (8–10) | III + 8.65 ± 0.49 (III 8–9) |
| JCC | 33.15 ± 0.35 (32–34) | 7.5 ± 0.42 (6–8) | 7.6 ± 0.31(5–7) | III + 19.35 ± 0.86 (III 18–20) | 9.05 ± 0.75 (8–10) | III + 6.45 ± 0.31 (III 6–7) |
| 3nJB | 33.45 ± 0.74(32–34) | 7.7 ± 0.51 (7–8) | 7.45 ± 0.60 (7–9) | III + 17.25 ± 1.07 (III 15–20) | 7.65 ± 0.67 (6–9) | III + 7.20 ± 0.61 (III 6–8) |
| 4nJB | 32.05 ± 0.22(32–33) | 7.1 ± 0.31 (7–8) | 7.95 ± 0.22 (7–8) | III +17.35 ± 0.49 (III 17–18) | 8.70 ± 0.47 (8–9) | III +7.15 ± 0.37 (III 7–8) |

Histological sectioning was used to examine gonad development in 4nJB and 3nJB. The testes of the 15-month-old 4nJB were at stage IV, in which a number of secondary spermatocytes were observed in the seminiferous tubules (Figure 1 G, the right). The ovaries of the 15-month-old 4nJB were at stage II, were rich in oocytes in synchronized development and were characterized by the location of the yolk nucleus near the cell nucleus (Figure 1 G, the left). In addition, during the reproductive season, the water-like semen and the mature eggs could be stripped out from the two-year-old male and female 4nJB individuals, respectively. The male 4nJB and female 4nJB were used for self-mating and viable F_2_ hybrids (details not shown) were produced. These results suggested that 4nJB are fertile, but at a reduced rate compared with their parents. By contrast, there were three types of gonadal structure in the triploid hybrids. The first type was testis-like gonads that comprised many lobules containing numerous spermatides. Some degenerated spermatids were found and no mature spermatozoa were observed (Figure 1 H, the right). The second type was ovary-like gonads comprising many nests of small, undeveloped cells and a few small growing oocytes, as well as enlarged and degenerated oocytes (Figure 1 H, the left). The third type only had fat tissue where the gonads should have been and neither testes nor ovaries were observed. In the reproductive season, no milt or eggs could be stripped out from the one and two-year old males and females of 3nJB. These results suggest that the 3nJB hybrids are sterile.

### Liver transcriptome analysis

cDNA libraries were prepared from liver RNA samples from 3nJB, 4nJB and both parents (JCC and BSB). The libraries were subjected to next-generation sequencing and the sequencing reads were processed to obtain clean reads. The clean reads for these libraries have been uploaded to the NCBI Sequence Read Archive site (http://www.ncbi.nlm.nih.gov/sra/; accession nos. SRX1999729, SRX685580, SRX1998501 and SRX1999073.

We identified 4740 orthologous genes (Table S2) among the transcriptomes of 3nJB, 4nJB and their parents. Based on these orthologous genes and the distribution of cDNA variations, chimeric gene patterns were identified in 4nJB and 3nJB. We classified 10 gene patterns within three categories in 3nJB and 4nJB (Figure 2, Table 7). The first category includes patterns 1 and 2 (Figure 2 A and B), in which the genes are not chimeras or offspring-specific variations but are derived exclusively from one parent. Patterns 1 represents a gene from JCC and patterns 2 represents a gene from BSB. The first category comprised 44.50% and 49.86% of genes in the overlapping mapped regions in 4nJB and 3nJB, respectively. The second category includes patterns 3–8 (Figure 2 C–H, respectively), which are chimeric genes with offspring-specific variations, such as DNA insertions, DNA deletions or locus mutations. Patterns 3 and 4 represent a gene from JCC and BSB, respectively, but with locus mutations; patterns 5 and 6 represent a gene from JCC and BSB, respectively, but with DNA deletions; patterns 7 and 8 represent a gene from JCC and BSB, respectively, but with DNA insertions. The second category comprised 23.82% and 17.82% of genes in overlapping mapped regions in 4nJB and 3nJB, respectively. The third category includes patterns 9–10 (Figure 2 I and J), where chimeric genes have single or multiple fragments comprising continuous, alternating variations from parents-specific variants, with or without offspring-specific variations. Patterns 9 represent a gene from JCC, but with single or multiple fragments comprising continuous, alternating variations from BSB. Patterns 10 represent a gene from BSB, but with single or multiple fragments comprising continuous, alternating variations from JCC. The third category comprised 31.68% and 32.32% of genes in overlapping mapped regions in 4nJB and 3nJB, respectively.

**Table 7.**
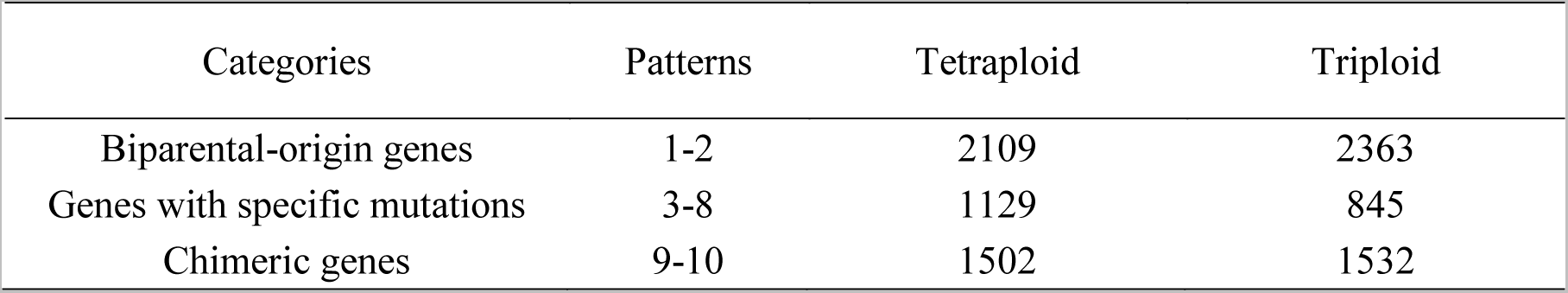
Gene numbers of each variation pattern in tetraploid and triploid using Japanese crucian carp as reference

**Figure 2.**
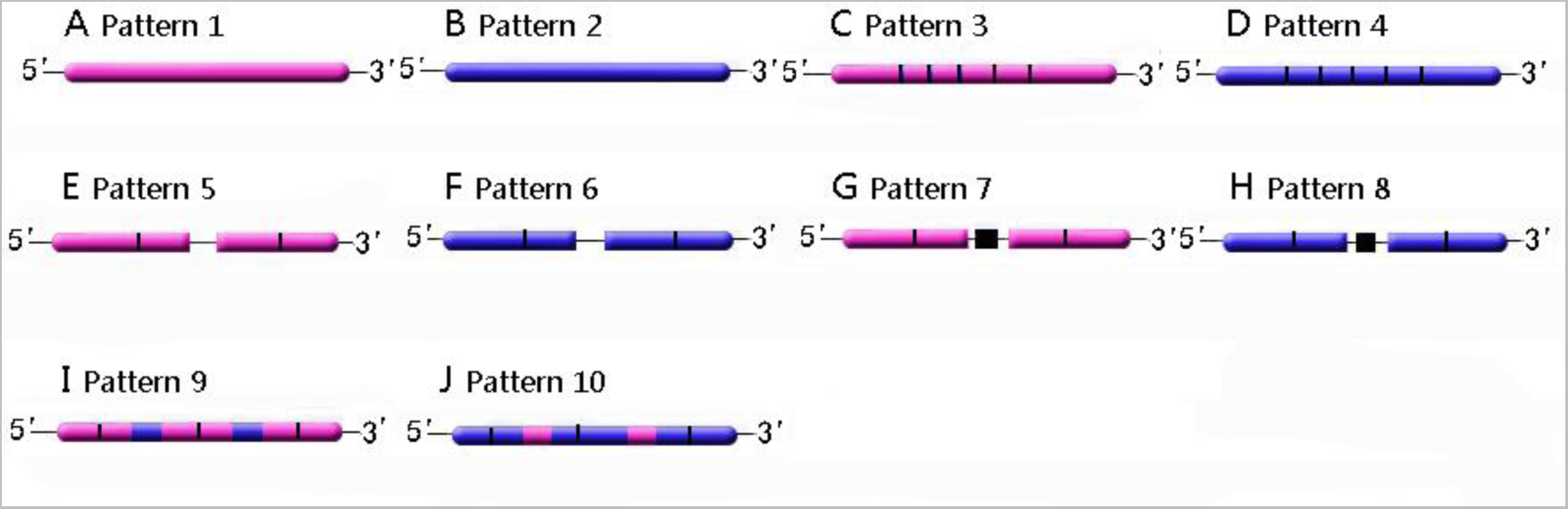
Schematic diagrams of gene patterns for 3nJB and 4nJB from hybridizing JCC (J) and BSB (B). Magenta bars marked J-variation denote offspring fragments with JCC-specific variants, blue bars marked B-variation show BSB-specific variants, and black bars marked F-variation show offspring-specific variants. Genes were classified as three categories. The first category include patterns 1–2 (A and B), which are not chimeras and offspring-specific variations and the genes are derived exclusively from one parent. The second category includes patterns 3–8(C–H, respectively) in which genes with offspring-specific variations, such as DNA insertions or the DNA deletions or the DNA locus mutations. The third category includes patterns 9–10 (I and J) where chimeric genes have single or multiple fragments consisting of continuous, alternating variations from parents-specific variants, and with or without offspring-specific variations.

### Chimeric genes validated in parental and hybrid fishes

The altered gene patterns were confirmed at the genomic level by PCR amplification and Sanger sequencing. Sanger sequencing validated 22 of the tested 30 chimeric genes, suggesting that the bioinformatics analysis identified 73.3% of chimeric genes correctly). For example, both *Basic helix-loop-helix family member e41*(*bh1he41*) (Figure S1) and *TATA-box binding protein associated factor 8*(*taf8*) (Figure S2) are exclusively from JCC in 4nJB; both *Phosphatidylinositol binding clathrin assembly protein* (*picalma*) (Figure S3) and *RNA binding motif protein 19* (*rbm19*) (Figure S4) are exclusively from JCC in 3nJB; *A-Raf proto-oncogene, serine/threonine kinase (araf)* (Figure S5), *SEC24 homolog C, COPII coat complex component*(*sec24c*) (Figure S6), *Chondroitin polymerizing factor a (chpfa)* (Figure S7), *SNAP associated protein*(*snapin*) (Figure S8) and *Tuberous sclerosis 1a* (*tsc1a*) (Figure S9) are exclusively from BSB in 3nJB; *Tripartite motif containing 71*(*trim71*) (Figure S10) and *RAN binding protein 9* (*ranbp9*) (Figure S11), have some DNA locus mutations in both 3nJB and 4nJB; *Phosphatidylinositol binding clathrin assembly protein* (*picalma*) (Figure S3) has three bases deleted in 4nJB; *Hexamethylene bis-acetamide inducible 1* (*hexim1*) (Figure S12) also has some base deleted in 4nJB; *SET translocation (myeloid leukemia-associated) B (setb)* (Figure S13) has three base inserted in 4nJB; *Tetratricopeptide repeat domain 4* (*ttc4*) (Figure S14) and *Immediate early response 5* (*ier5*) (Figure S15) also have some base inserted in 4nJB; *Pleckstrin homology-like domain, family B, member 2a* (*phldb2a*) (Figure S16) has one fragment derived from BSB and other fragments derived from JCC both in 3nJB and 4nJB; *CD2-associated protein* (*cd2ap*) (Figure S17) has one fragment derived from BSB and other fragments derived from JCC in 4nJB; *Mahogunin, ring finger 1b* (*mgrn1b*) (Figure S18) has one fragment derived from BSB and other fragments derived from JCC in 3nJB; *Beta-carotene oxygenase 2a* (*bco2a*) (Figure S19) has one fragment derived from JCC and other fragments derived from BSB both in 3nJB and 4nJB; *Kruppel-like factor 6a* (*klf6a*) (Figure S20) and *SR-related CTD-associated factor 1*(*scaf1*) (Figure S21) has one fragment derived from JCC and other fragments derived from BSB in 4nJB. *Calcium/calmodulin-dependent protein kinase II gamma* (*camk2g*) (Figure S22) has one fragment derived from JCC and other fragments derived from BSB in 3nJB;

**Figure 3.**
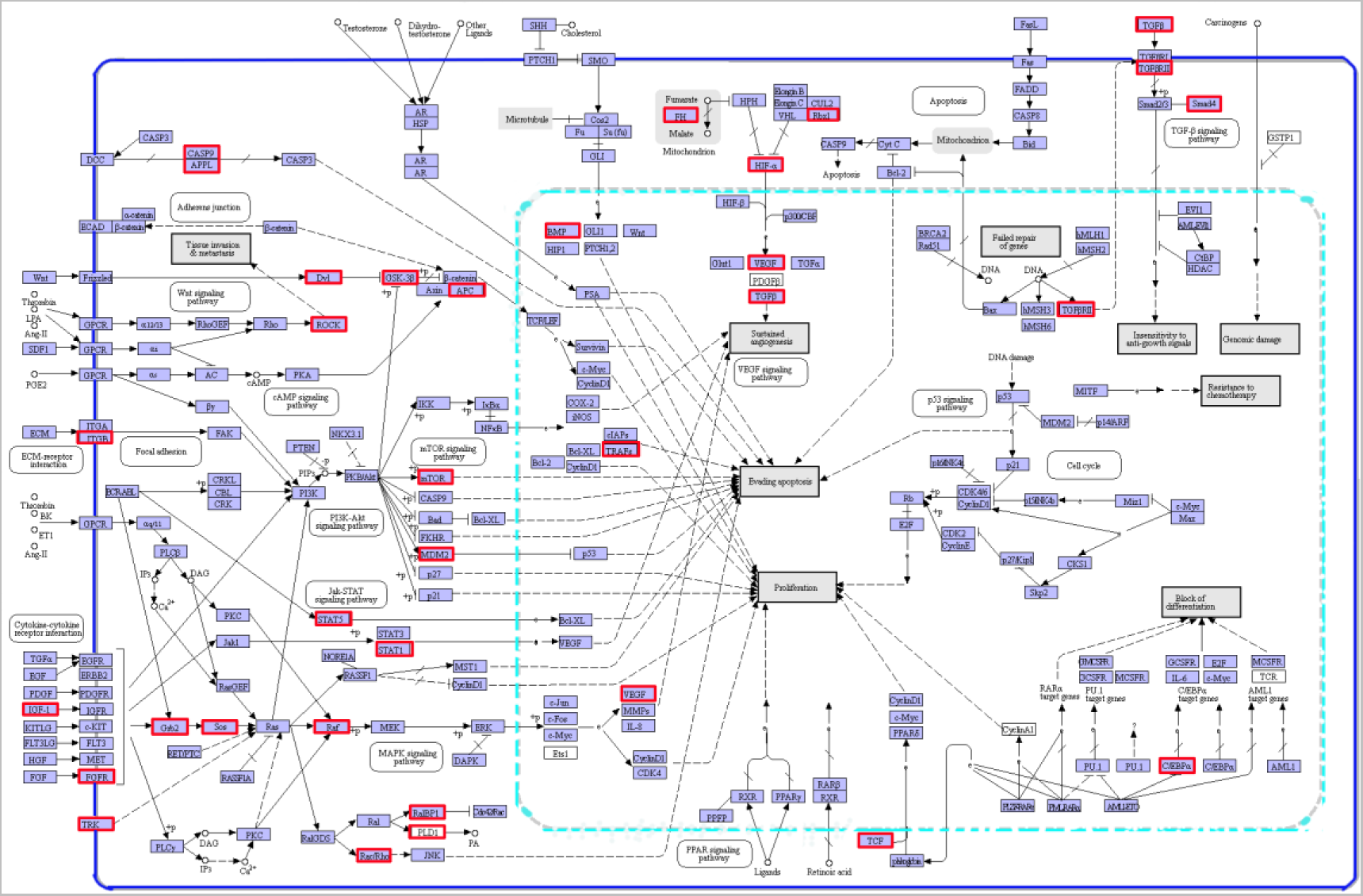
Representative chimeric genes involved pathways (map modified from KEGG pathway ko05200). Genes in Red frames are chimeric genes detected in allopolyploid hybrids. Pathways in dark blue frames are chimeric genes involved pathways.

The sequences of 22 homologous genes for 4nJB and 3nJB are available at NCBI GenBank (Table S3). Moreover, we mapped these validated chimeric genes from the 4nJB and 3nJB transcriptomes to the homologous genes in the BSB reference genome to determine the structural changes (S1–22_Fig). PCR analysis *kcnk5b* showed that *kcnk5b* has some missense mutations and 12 bp insertion in 4nJB (Figure 4).

**Figure 4.**
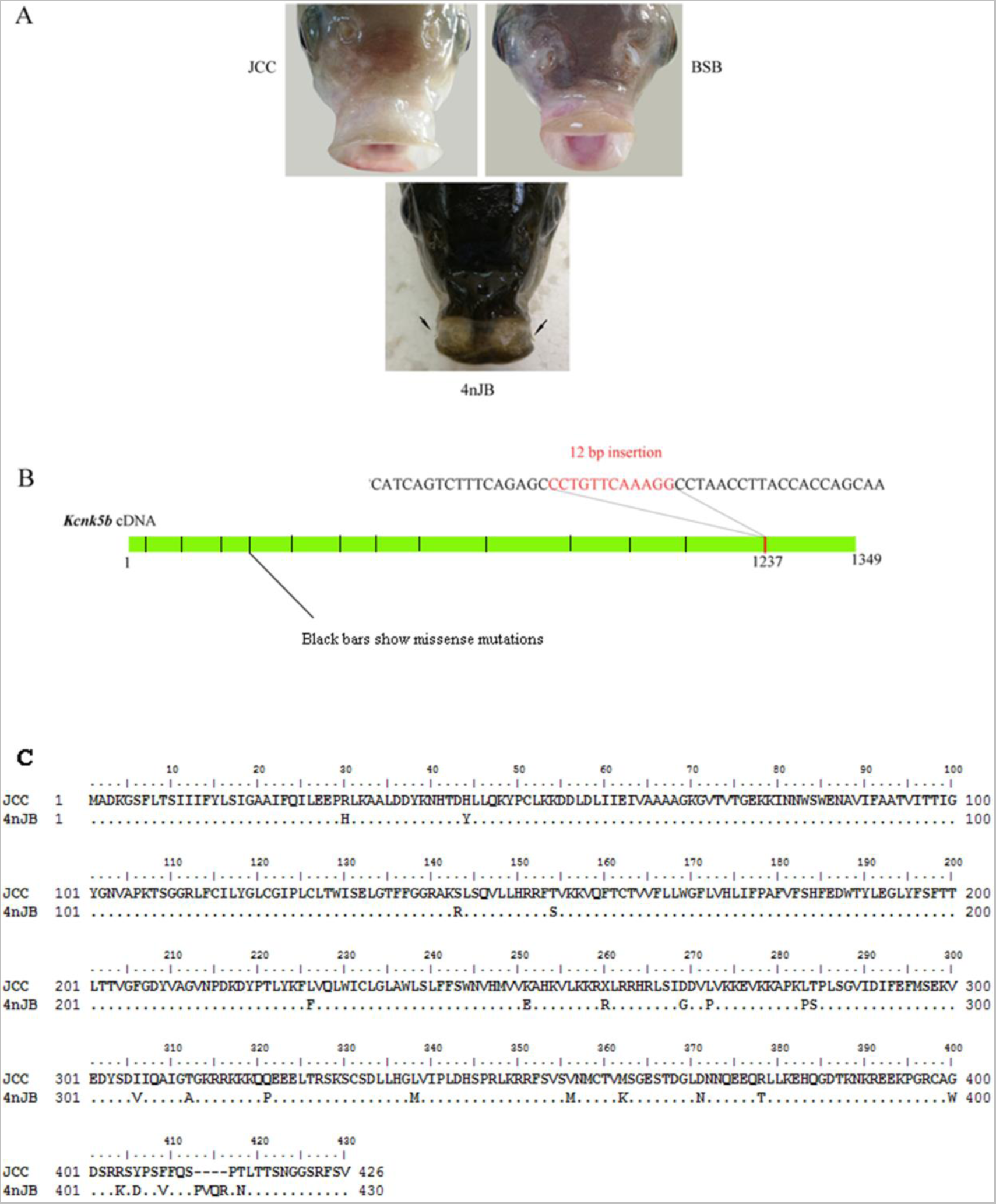
The 4nJB phenotype is related to gain-of-function mutations within the K^+^ channel *kcnk5b*. (A) The 4nJB had a pair of short barbels while there were no barbels in their parents (B) 4nJB harbor an intragenic insertion and missense mutations in *kcnk5b*. (C) The amino acids variation in the 4nJB compared with JCC.

## Disscussion

Few hybridization and allopolyploidization studies have focused on animals. This study provided new insights into allopolyploidization related genomic variation in vertebrates and a preliminary genomic characterization of 4nJB and 3nJB, revealing significant insights into the evolution and genome function of allopolyploid animals.

### The formation mechanism of allopolyploid fish

Distant hybridization is a useful approach to produce polyploid hybrid lineages in plants and animals. However, much remains unknown about the processes and consequences of allopolyploidization[40]. Previous studies have concluded that distant hybridization is more likely to form diploid or triploid hybrids in the first generation when the parents have the same chromosome number. However, there have been few reports that tetraploid and triploid hybrids are formed in the first generation of distant hybridization when the parents have different chromosome numbers [2, 24].

In general, allopolyploid formation can result from three events: the first is somatic chromosome doubling in a diploid hybrid; the second is the fusion of two unreduced gametes after failure of reduction divisions in meiosis; and the third is unreduced (diploid) gametes fusion with normal haploid gametes to form triploids [3, 41, 42]. In this study, both tetraploid and triploid hybrids were produced by crossing female Japanese crucian carp and male blunt snout bream. Somatic chromosome doubling after fertilization is the most likely explanation for the formation 4nJB (Figure 5, right). Somatic chromosome doubling in certain creatures is related to a failure of cell division following mitotic doubling. It may arise in the zygote, early embryo, or meristem of a plant, and will ultimately results the formation of polyploidy tissues and polyploids species [43, 44]. The naturally occurring tetraploid among Lamarckian primrose was produced by zygote chromosome doubling [45]. The 3nJB formation probably occurred because second polar body extrusion was inhibited during the second division of meiosis (Figure 5, left). It plays an important role that the sister chromatids do not separate or the second polar body cannot release normally in the second meiotic division. Triploids appeared in the descendants of the open-pollinated diploid *Crepis capillaris*, and they appeared to have been produced by the fusion of reduced haploid gametes and unreduced diploid gametes [44] Both allotetraploid and allotriploid have to experience genomic changes for surviving.

**Figure 5.**
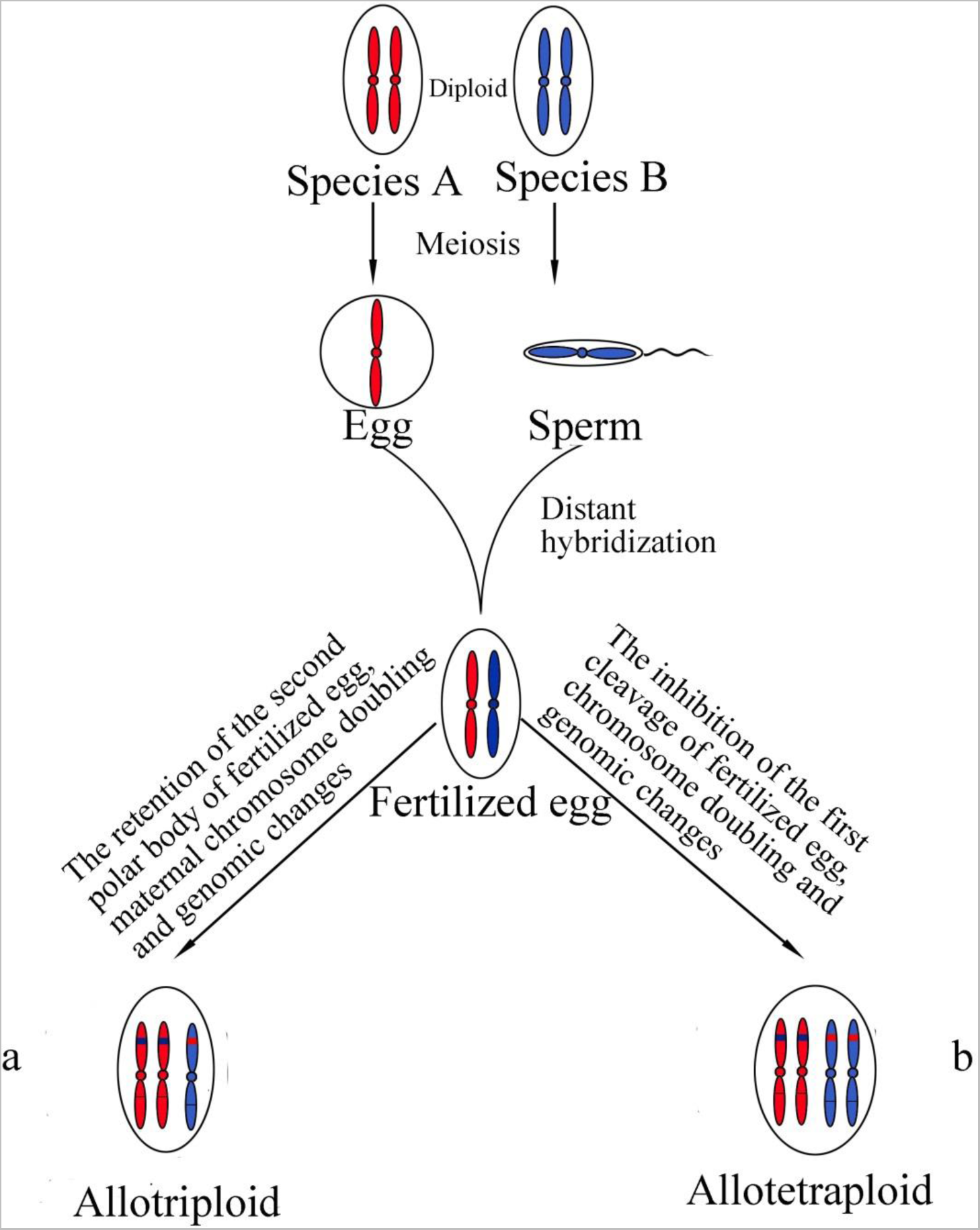
Illustration of allopolyploids. For simplicity, only one pair of homologous chromosomes (either in red or blue) is shown in a diploid. (a) Formation of an allotriploid by doubling maternal chromosomes. (b) Formation of an allotetraploid by interspecific hybridization followed by chromosome doubling. Both allotetraploid and allotriploid have to experience genomic changes for surviving.

### The related mechanisms of high frequency chimeric and mutated genes

The survival of allopolyploid offspring may be driven severely by the potential comprehensive effects related to genomic structural change. Previous studies showed that the genomes of newly formed natural or artificial polyploids may experience rapid gene loss, genome restructuring, and altered patterns of gene expression [46–50]. Polyploids of Tragopogon [51, 52], Brassica [10, 53], and wheat [54, 55], exhibit relatively high levels of genomic rearrangements.

The tetraploid and triploid hybrids reported here, which were produced from different subfamilies with the JCC and BSB genomes, indicated a “genetic melting” of the two organisms. We identified patterns of variation among orthologous genes in the newly formed hybrids, and demonstrated high proportions of chimeric and mutated genes in the hybrid genomes. The main effect of distant hybridization is to cause genetic variation in the hybrid offspring [2]. JCC and BSB belong to different subfamily and have different number chromosome. Using them as parents for the hybridization, it increases nuclear-nuclear and nuclear-cytoplasmic incompatibility between the two different species, which easily gives rise to drastic genomic imbalances in allopolyploid hybrids [56].

The high frequency of chimeric and mutated genes may be driven by erroneous DNA excision between homologous parental genes [57]. In allopolyploids, genome changes (deletions, duplications, and translocations) are usually caused by large-scale DNA repair via recombination, nonhomologous end-joining or transposon activity [58–61]. Studies in *Brassica napus* polyploids indicated that homoeologous pairing recombination is a key mechanism for genome restructuring [62, 63]. Genetic recombination also has been reported in allopolyploid clawed frogs [64] and salamanders [65]. Abnormalities of DNA or RNA repair, such as dysfunction of RAD or other genes and pathways, may contribute to the high rate of homologous recombination in the polyploidy hybrids [61, 66]. Both in 4nJB and 3nJB, chimeric genes and mutated genes were involved directly involved in the regulation of cell cycle and DNA damage response and repair (Figure 3).

The 4nJB and 3nJB contain combinations of the JCC and BSB genomes in a common nucleus, which might have allowed homologous chromosomes reciprocal exchange and gene conversion to occur more easily, leading to large genetic changes in the newly formed 4nJB and 3nJB. The common occurrence of chimeric genes and mutated genes that both exists in two different ploidies hybrid fish might result from different processes that relate to chimeras: imprecise excision of an unpaired duplication during large-loop mismatch repair or replication slippage [67]. Chimeric genes and mutational genes might have formed after the two different genomes merged and before the whole genome duplication that leads to allopolyploidization [67]. Recent reports of higher mutation rates in heterozygotes support this possibility.

Previous studies showed that repetitive elements (including transposable elements and tandem repeats) play an irreplaceable role in generating new chimerical genes, which flanking the paralogs can mediate recombinations to form chimeras [68]. Hybridization and allopolyploidization might trigger extensive transposon activity, which can result in extensive chimeric regions [49, 60, 69–71]. Our previous study showed that some new transposons could burst allopolyploid hybrids genomic and driving for genomic evolution [72]. Some transposable elements are capable of mobilizing adjacent sequence, thus new chimeric genes were generated [68, 73, 74]. In additional, during the process of DNA replication, double-strand break (DSB) is more likely occur at tandem repeats locus and then the chimera is formed through recombination between different sequences in the process of DSB repair. Our analyses cannot identify the exact mechanism.

### The morphological and physiological differences related to chimeric genes and mutated genes

At the molecular level, we still know relatively little about diverse genetic mechanisms underlie novel phenotypes traits in hybrids. The hypothesis that particular genetic mechanisms influence the outcome of hybridization via their effect on phenotypes has been tested severely in very few systems [75–78]. Structural variation of hybrids offspring, including chromosomal rearrangements, gene insertion or loss and transposable element distribution, can produce substantial morphological and physiological effects and directly impact recombination rate and reproductive compatibility with their parental species [79, 80]. The differences of parental genome structure can induce further restructuring after recombination of the hybrid genomes [[58]. Both chimeric genes and mutated genes might produce gene structural changes that would decrease the activity of an encoded enzyme activity or its fidelity, by affecting normal transcriptional processing, thus may relate to morphological and physiological changes in the hybrids [81].

The phenotypic characteristics of 4nJB (Figure 1C) and 3nJB (Figure 1D) showed obvious differences compared with JCC(Figure 1A) and BSB (Figure 1B). The most obvious variation was that the 4nJB had barbels while there were no barbels in the 3nJB hybrids or their parents (Figure 4A). Studied zebrafish *another longfin* (*alf*) mutant showed that *potassium channel, subfamily K, member 5b* (*kcnk5b*) mutations lead to proportionally enlarged fins and barbels [82]. In this study, we found there is some missense mutation and an intragenic insertion in the K+ channel *kcnk5b*, which may be related to the novel presence of the barbels in 4nJB (Figure 4 B and C). The genomic changes in 4nJB led to the spontaneous development of barbels, which is of huge evolutionary significance. In terms of fertility, the two types of hybrids showed different characteristics. Histological observation showed that the 4nJB had normal gonadal development (Figure 1, G and H) and produced mature sperm and eggs. Further self-mating experiments proved that the 4nJB had reduced fertility. However, the gonadal development of 3nJB was retarded, asynchronous and exhibited abundant polymorphisms. Genetic recombination not only generates novel gene combinations and phenotypes, but also might damage the karyotype and lead to aberrant meiotic behavior and reduced fertility [58]. The 4nJB have reduced fertility, which might have been caused by aberrant meiotic behavior. The 3nJB are completely infertile, which might have been caused by incompatible genomes and abnormal expression and regulation of genes related to gonad development [83, 84].

In addition, allopolyploidization occurs less frequently in vertebrates than in plants, possibly because of the severe effect of genome shock, leading to greatly reduced viability of F_1_ hybrid offspring [6–8]. In this study, the two different ploidy allopolyploid hybrid fish were produced by same parents; however, the proportion of variant orthologous genes in 4nJB (54.3%) was higher than in 3nJB (49.1%). The result showed that although the genomes of two types of hybrids shared the same origin, they experienced different degrees of variation. On the basis of the chromosomal number and karyotypes of 4nJB and 3nJB, 4nJB appears to have two sets of chromosomes from JCC and two sets from BSB, and 3nJB appears to have two sets of chromosomes from JCC and one set from BSB. We speculated that mutations and recombination of the genomes of the 4nJB hybrids were easier to induce because the different combination of chromosome numbers caused different variations in the genome.

## Supporting information

**Tables SI**

Table S1. Primers sequences used in the PCR.

Table S2. Information on the homologous genes from allotetraploid hybrids (4nJB), allotriploid hybrids (3nJB), Japanese crucian carp (JCC) and blunt snout bream (BSB).

Table S3. The orthologous genes among the transcriptomes of 3nJB, 4nJB and their parents (JCC and BSB).

**Figures SI**

Figure S1. Diagram and alignment of bhlhe41 from Japanese crucian carp (JCC), blunt snout bream (BSB), tetraploid(4nJB) and triploid (3nJB). (A): Schematic map of relationships between the reference genome, Sanger sequences and contigs of the de novo assembly. (B): Sequence alignment for bhlhe41.

Figure S2. Diagram and alignment of taf8 from Japanese crucian carp (JCC), blunt snout bream (BSB), triploid hybrids (3nJB) and tetraploid hybrids (4nJB). (A): Schematic map of relationships between the reference genome, Sanger sequences and contigs of the de novo assembly. (B): Sequence alignment for taf8.

Figure S3. Diagram and alignment of picalma from Japanese crucian carp (JCC), blunt snout bream (BSB), tetraploid hybrids (4nJB) and triploid hybrids (3nJB). (A): Schematic map of relationships between the reference genome, Sanger sequences and contigs of the de novo assembly. (B): Sequence alignment for picalma.

Figure S4. Diagram and alignment of rbm19 from Japanese crucian carp (JCC), blunt snout bream (BSB), tetraploid hybrids (4nJB) and triploid hybrids (3nJB). (A): Schematic map of relationships between the reference genome, Sanger sequences and contigs of the de novo assembly. (B): Sequence alignment for rbm19.

Figure S5. Diagram and alignment of araf from Japanese crucian carp (JCC), blunt snout bream (BSB), tetraploid hybrids (4nJB) and triploid hybrids (3nJB). (A): Schematic map of relationships between the reference genome, Sanger sequences and contigs of the de novo assembly. (B): Sequence alignment for araf.

Figure S6. Diagram and alignment of sec24c from Japanese crucian carp (JCC), blunt snout bream (BSB), tetraploid hybrids (4nJB) and triploid hybrids (3nJB). (A): Schematic map of relationships between the reference genome, Sanger sequences and contigs of the de novo assembly. (B): Sequence alignment for sec24c.

Figure S7. Diagram and alignment of chpfa from Japanese crucian carp (JCC), blunt snout bream (BSB), triploid hybrids (3nJB) and tetraploid hybrids (4nJB). (A): Schematic map of relationships between the reference genome, Sanger sequences and contigs of the de novo assembly. (B): Sequence alignment for chpfa.

Figure S8 Diagram and alignment of snapin from Japanese crucian carp (JCC), blunt snout bream (BSB), triploid hybrids (3nJB) and tetraploid hybrids (4nJB). (A): Schematic map of relationships between the reference genome, Sanger sequences and contigs of the de novo assembly. (B): Sequence alignment for snapin.

Figure S9. Diagram and alignment of tasc1a from Japanese crucian carp (JCC), blunt snout bream (BSB), triploid hybrids (3nJB) and tetraploid hybrids (4nJB). (A): Schematic map of relationships between the reference genome, Sanger sequences and contigs of the de novo assembly. (B): Sequence alignment for tasc1a.

Figure S10. Diagram and alignment of trim71 from Japanese crucian carp (JCC), blunt snout bream (BSB), tetraploid hybrids (4nJB) and triploid hybrids (3nJB). (A): Schematic map of relationships between the reference genome, Sanger sequences and contigs of the de novo assembly. (B): Sequence alignment for trim71.

Figure S11. Diagram and alignment of ranbp9 from Japanese crucian carp (JCC), blunt snout bream (BSB), tetraploid hybrids (4nJB) and triploid hybrids (3nJB). (A): Schematic map of relationships between the reference genome, Sanger sequences and contigs of the de novo assembly. (B): Sequence alignment for ranbp9.

Figure S12. Diagram and alignment of hexim1 from Japanese crucian carp (JCC), blunt snout bream (BSB), triploid hybrids (3nJB) and tetraploid hybrids (4nJB). (A): Schematic map of relationships between the reference genome, Sanger sequences and contigs of the de novo assembly. (B): Sequence alignment for hexim1.

Figure S13 Diagram and alignment of setb from Japanese crucian carp (JCC), blunt snout bream (BSB), tetraploid hybrids (4nJB) and triploid hybrids (3nJB). (A): Schematic map of relationships between the reference genome, Sanger sequences and contigs of the de novo assembly. (B): Sequence alignment for setb.

Figure S14 Diagram and alignment of ttc4 from Japanese crucian carp (JCC), blunt snout bream (BSB), tetraploid hybrids (4nJB) and triploid hybrids (3nJB). (A): Schematic map of relationships between the reference genome, Sanger sequences and contigs of the de novo assembly. (B): Sequence alignment for ttc4.

Figure S15. Diagram and alignment of ier5 from Japanese crucian carp (JCC), blunt snout bream (BSB), triploid hybrids (3nJB) and tetraploid hybrids (4nJB). (A): Schematic map of relationships between the reference genome, Sanger sequences and contigs of the de novo assembly. (B): Sequence alignment for ier5.

Figure S16. Diagram and alignment of phldb2a from Japanese crucian carp (JCC), blunt snout bream (BSB), tetraploid hybrids (4nJB) and triploid hybrids (3nJB). (A): Schematic map of relationships between the reference genome, Sanger sequences and contigs of the de novo assembly. (B): Sequence alignment for phldb2a.

Figure S17. Diagram and alignment of cd2ap from Japanese crucian carp (JCC), blunt snout bream (BSB) and tetraploid hybrids (4nJB). (A): Schematic map of relationships between the reference genome, Sanger sequences and contigs of the de novo assembly. (B): Sequence alignment for cd2ap.

Figure S18. Diagram and alignment of mgrn1b from Japanese crucian carp (JCC), blunt snout bream (BSB), triploid hybrids (3nJB) and tetraploid hybrids (4nJB). (A): Schematic map of relationships between the reference genome, Sanger sequences and contigs of the de novo assembly. (B): Sequence alignment for mgrn1b.

Figure S19. Diagram and alignment of bco2a from Japanese crucian carp (JCC), blunt snout bream (BSB), tetraploid hybrids (4nJB) and triploid hybrids (3nJB). (A): Schematic map of relationships between the reference genome, Sanger sequences and contigs of the de novo assembly. (B): Sequence alignment for bco2a.

Figure S20. Diagram and alignment of klf6a from Japanese crucian carp (JCC), blunt snout bream (BSB) and tetraploid hybrids (4nJB). (A): Schematic map of relationships between the reference genome, Sanger sequences and contigs of the de novo assembly. (B): Sequence alignment for klf6a.

Figure S21. Diagram and alignment of scaf1 from Japanese crucian carp (JCC), blunt snout bream (BSB) and tetraploid hybrids (4nJB). (A): Schematic map of relationships between the reference genome, Sanger sequences and contigs of the de novo assembly. (B): Sequence alignment for scaf1.

Figure S22. Diagram and alignment of camk2g from Japanese crucian carp (JCC), blunt snout bream (BSB), tetraploid hybrids (4nJB) and triploid hybrids (3nJB). (A): Schematic map of relationships between the reference genome, Sanger sequences and contigs of the de novo assembly. (B): Sequence alignment for camk2g.

## Acknowledgement

This research was supported by National Natural Science Foundation of China Grants 30930071, 91331105, 31360514, 31430088, and 31210103918, the Cooperative Innovation Center of Engineering and New Products for Developmental Biology of Hunan Province (20134486), the Construction Project of Key Discipline of Hunan Province and China, the National High Technology Research and Development Program of China (Grant No. 2011AA100403).

## References

1. Biradar, D., and Rayburn, A.L. (1993). Heterosis and nuclear DNA content in maize. HEREDITY-LONDON- 71, 300–300.

2. Liu, S. (2010). Distant hybridization leads to different ploidy fishes. Science China Life Sciences 53, 416–425.

3. Mallet, J. (2007). Hybrid speciation. Nature 446, 279–283.

4. Bullini, L. (1994). Origin and evolution of animal hybrid species. Trends in ecology & evolution 9, 422–426.

5. Muller, H. (1925). Why polyploidy is rarer in animals than in plants. The American Naturalist 59, 346–353.

6. Orr, H.A. (1990). "Why Polyploidy is Rarer in Animals Than in Plants" Revisited. American Naturalist, 759–770.

7. Mable, B. (2004). ‘Why polyploidy is rarer in animals than in plants’: myths and mechanisms. Biological Journal of the Linnean Society 82, 453–466.

8. Wertheim, B., Beukeboom, L., and Van de Zande, L. (2013). Polyploidy in animals: effects of gene expression on sex determination, evolution and ecology. Cytogenetic and genome research 140, 256–269.

9. Song, K., Lu, P., Tang, K., and Osborn, T.C. (1995). Rapid genome change in synthetic polyploids of Brassica and its implications for polyploid evolution. Proceedings of the National Academy of Sciences 92, 7719–7723.

10. Gaeta, R.T., Pires, J.C., Iniguez-Luy, F., Leon, E., and Osborn, T.C. (2007). Genomic changes in resynthesized Brassica napus and their effect on gene expression and phenotype. The Plant Cell 19, 3403–3417.

11. Otto, S.P., and Whitton, J. (2000). Polyploid incidence and evolution. Annual review of genetics 34, 401–437.

12. Yu, X. (1989). China freshwater fisheries chromosome. Science.

13. Anamthawat-JÓnsson, K., Humphreys, M., and Joness, R. (1997). Novel diploids following chromosome elimination and somatic recombination in Lolium multiflorum× Festuca arundinacea hybrids. Heredity 78.

14. Kenton, A., Parokonny, A.S., Gleba, Y.Y., and Bennett, M.D. (1993). Characterization of the Nicotiana tabacum L. genome by molecular cytogenetics. Molecular and General Genetics MGG 240, 159–169.

15. Wendel, J.F. (2000). Genome evolution in polyploids. In Plant molecular evolution. (Springer), pp. 225–249.

16. Ozkan, H., Levy, A.A., and Feldman, M. (2001). Allopolyploidy-induced rapid genome evolution in the wheat (Aegilops–Triticum) group. The Plant Cell 13, 1735–1747.

17. Galili, G., Levy, A.A., and Feldman, M. (1986). Gene-dosage compensation of endosperm proteins in hexaploid wheat Triticum aestivum. Proceedings of the National Academy of Sciences 83, 6524–6528.

18. Arnheim, N., Krystal, M., Schmickel, R., Wilson, G., Ryder, O., and Zimmer, E. (1980). Molecular evidence for genetic exchanges among ribosomal genes on nonhomologous chromosomes in man and apes. Proceedings of the National Academy of Sciences 77, 7323–7327.

19. Hu, W., Timmermans, M.C., and Messing, J. (1998). Interchromosomal recombination in Zea mays. Genetics 150, 1229–1237.

20. Soltis, D.E., and Soltis, P.S. (1995). The dynamic nature of polyploid genomes. Proceedings of the National Academy of Sciences 92, 8089–8091.

21. Liu, B., and Wendel, J.F. (2002). Non-Mendelian phenomena in allopolyploid genome evolution. Current Genomics 3, 489–505.

22. Liu, S., Liu, Y., Zhou, G., Zhang, X., Luo, C., Feng, H., He, X., Zhu, G., and Yang, H. (2001). The formation of tetraploid stocks of red crucian carp× common carp hybrids as an effect of interspecific hybridization. Aquaculture 192, 171–186.

23. Wolters W R, C.C.L., Libey G S. (1982). Erythrocyte nuclear measurements of diploid and triploid channel catfish, Ictalurus punctatus (Rafinesque)[J]. Journal of Fish Biology 20, 253–258.

24. Xiao, J., Kang, X., Xie, L., Qin, Q., He, Z., Hu, F., Zhang, C., Zhao, R., Wang, J., and Luo, K. (2014). The fertility of the hybrid lineage derived from female Megalobrama amblycephala× male Culter alburnus. Animal reproduction science 151, 61–70.

25. Liu, Y. (1993). Propagation physiology of main cultivated fish in China. Beijing: Agricultural Publishing House 147, 147–148.

26. Dion-Côté, A.-M., Renaut, S., Normandeau, E., and Bernatchez, L. (2014). RNA-seq reveals transcriptomic shock involving transposable elements reactivation in hybrids of young lake whitefish species. Molecular biology and evolution 31, 1188–1199.

27. Anders, S., and Huber, W. (2010). Differential expression analysis for sequence count data. Genome biol 11, R106.

28. Reeb, P.D., and Steibel, J.P. (2013). Evaluating statistical analysis models for RNA sequencing experiments. Frontiers in genetics 4.

29. Flicek, P., Ahmed, I., Amode, M.R., Barrell, D., Beal, K., Brent, S., Carvalho-Silva, D., Clapham, P., Coates, G., Fairley, S., et al. (2013). Ensembl 2013. Nucleic acids research 41, D48–55.

30. Ye, J., Fang, L., Zheng, H., Zhang, Y., Chen, J., Zhang, Z., Wang, J., Li, S., Li, R., Bolund, L., et al. (2006). WEGO: a web tool for plotting GO annotations. Nucleic acids research 34, W293–297.

31. Blanc, G., and Wolfe, K.H. (2004). Widespread paleopolyploidy in model plant species inferred from age distributions of duplicate genes. The Plant Cell 16, 1667–1678.

32. Hall, T.A. (1999). BioEdit: a user-friendly biological sequence alignment editor and analysis program for Windows 95/98/NT. In Nucleic acids symposium series, Volume 41. pp. 95–98.

33. Li, H., and Durbin, R. (2009). Fast and accurate short read alignment with Burrows-Wheeler transform. Bioinformatics 25, 1754–1760.

34. Li, H., Handsaker, B., Wysoker, A., Fennell, T., Ruan, J., Homer, N., Marth, G., Abecasis, G., and Durbin, R. (2009). The Sequence Alignment/Map format and SAMtools. Bioinformatics 25, 2078–2079.

35. Van der Auwera, G.A., Carneiro, M.O., Hartl, C., Poplin, R., Del Angel, G., Levy-Moonshine, A., Jordan, T., Shakir, K., Roazen, D., Thibault, J., et al. (2013). From FastQ data to high confidence variant calls: the Genome Analysis Toolkit best practices pipeline. Current protocols in bioinformatics / editoral board, Andreas D. Baxevanis … [et al.] 43, 11.10.11–33.

36. DePristo M.A., Banks, E., Poplin, R., Garimella, K.V., Maguire, J.R., Hartl, C., Philippakis, A.A., del Angel, G., Rivas, M.A., Hanna, M., et al. (2011). A framework for variation discovery and genotyping using next-generation DNA sequencing data. Nature genetics 43, 491–498.

37. McKenna, A., Hanna, M., Banks, E., Sivachenko, A., Cibulskis, K., Kernytsky, A., Garimella, K., Altshuler, D., Gabriel, S., Daly, M., et al. (2010). The Genome Analysis Toolkit: a MapReduce framework for analyzing next-generation DNA sequencing data. Genome Res 20, 1297–1303.

38. Danecek, P., Auton, A., Abecasis, G., Albers, C.A., Banks, E., DePristo, M.A., Handsaker, R.E., Lunter, G., Marth, G.T., Sherry, S.T., et al. (2011). The variant call format and VCFtools. Bioinformatics 27, 2156–2158.

39. Sambrook, J., Fritsch, E.F., and Maniatis, T. (1989). Molecular cloning, Volume 2, (Cold spring harbor laboratory press New York).

40. Abbott, R., Albach, D., Ansell, S., Arntzen, J., Baird, S., Bierne, N., Boughman, J., Brelsford, A., Buerkle, C., and Buggs, R. (2013). Hybridization and speciation. Journal of Evolutionary Biology 26, 229–246.

41. Husband, B.C. (2000). Constraints on polyploid evolution: a test of the minority cytotype exclusion principle. Proceedings of the Royal Society of London B: Biological Sciences 267, 217–223.

42. Ramsey, J., and Schemske, D.W. (2002). Neopolyploidy in flowering plants. Annual review of ecology and systematics, 589–639.

43. Volff, J.N. (2005). Genome evolution and biodiversity in teleost fish. Heredity 94, 280–294.

44. And, J.R., and Schemske, D.W. (1998). Pathways, mechanisms, and rates of polyploid formation in flowering plants. Annual Review of Ecology & Systematics 29, 467–501.

45. Gates, R.R. (1909). The stature and chromosomes of Oenothera gigas, De Vries. Zeitschrift Für Induktive Abstammungs Und Vererbungslehre 3, 220–220.

46. Adams, K.L., and Wendel, J.F. (2005). Polyploidy and genome evolution in plants. Current opinion in plant biology 8, 135–141.

47. Xiong, Z., Gaeta, R.T., and Pires, J.C. (2011). Homoeologous shuffling and chromosome compensation maintain genome balance in resynthesized allopolyploid Brassica napus. Proceedings of the National Academy of Sciences 108, 7908–7913.

48. Hegarty, M.J., Abbott, R.J., and Hiscock, S.J.(2012). Allopolyploid speciation in action: The origins and evolution of Senecio cambrensis. In Polyploidy and Genome Evolution. (Springer), pp. 245–270.

49. Madlung, A., Tyagi, A.P., Watson, B., Jiang, H., Kagochi, T., Doerge, R.W., Martienssen, R., and Comai, L. (2005). Genomic changes in synthetic Arabidopsis polyploids. The Plant Journal 41, 221–230.

50. Schnable, J.C., Springer, N.M., and Freeling, M. (2011). Differentiation of the maize subgenomes by genome dominance and both ancient and ongoing gene loss. Proceedings of the National Academy of Sciences 108, 4069–4074.

51. Ownbey, M. (1950). Natural hybridization and amphiploidy in the genus Tragopogon. American Journal of Botany, 487–499.

52. Buggs, R.J., Chamala, S., Wu, W., Tate, J.A., Schnable, P.S., Soltis, D.E., Soltis, P.S., and Barbazuk, W.B. (2012). Rapid, repeated, and clustered loss of duplicate genes in allopolyploid plant populations of independent origin. Current Biology 22, 248–252.

53. Liu, S., Liu, Y., Yang, X., Tong, C., Edwards, D., Parkin, I.A., Zhao, M., Ma, J., Yu, J., and Huang, S. (2014). The Brassica oleracea genome reveals the asymmetrical evolution of polyploid genomes. Nature communications 5.

54. Brenchley, R., Spannagl, M.,Pfeifer, M., Barker, G.L., D’Amore, R., Allen, A.M., McKenzie, N., Kramer, M., Kerhornou, A., and Bolser, D. (2012). Analysis of the bread wheat genome using whole-genome shotgun sequencing. Nature 491, 705–710.

55. Feldman, M., Liu, B., Segal, G., Abbo, S., Levy, A.A., and Vega, J.M. (1997). Rapid elimination of low-copy DNA sequences in polyploid wheat: a possible mechanism for differentiation of homoeologous chromosomes. Genetics 147, 1381–1387.

56. Zhang, Z., Chen, J., Li, L., Tao, M., Zhang, C., Qin, Q., Xiao, J., Liu, Y., and Liu, S. (2014). Research advances in animal distant hybridization. Science China Life Sciences 57, 889–902.

57. Liu, S., Luo, J., Chai, J., Ren, L., Zhou, Y., Huang, F., Liu, X., Chen, Y., Zhang, C., and Tao, M. (2016). Genomic incompatibilities in the diploid and tetraploid offspring of the goldfish× common carp cross. Proceedings of the National Academy of Sciences, 201512955.

58. Gaeta, R.T., and Chris Pires, J. (2010). Homoeologous recombination in allopolyploids: the polyploid ratchet. New Phytologist 186, 18–28.

59. Wang, H.-C., Chou, W.-C., Shieh, S.-Y., and Shen, C.-Y. (2006). Ataxia telangiectasia mutated and checkpoint kinase 2 regulate BRCA1 to promote the fidelity of DNA end-joining. Cancer Research 66, 1391–1400.

60. Fedoroff, N.V. (2012). Transposable elements, epigenetics, and genome evolution. Science 338, 758–767.

61. Huang, J., Huen, M.S., Kim, H., Leung, C.C.Y., Glover, J.M., Yu, X., and Chen, J. (2009). RAD18 transmits DNA damage signalling to elicit homologous recombination repair. Nature cell biology 11, 592–603.

62. Felsenstein, J. (1974). The evolutionary advantage of recombination. Genetics 78, 737–756.

63. Muller, H.J. (1964). The relation of recombination to mutational advance. Mutation Research/Fundamental and Molecular Mechanisms of Mutagenesis 1, 2–9.

64. Evans, B.J., Kelley, D.B., Melnick, D.J., and Cannatella, D.C. (2005). Evolution of RAG-1 in polyploid clawed frogs. Molecular biology and evolution 22, 1193–1207.

65. Bi, K., Bogart, J.P., and Fu, J. (2008). The prevalence of genome replacement in unisexual salamanders of the genus Ambystoma (Amphibia, Caudata) revealed by nuclear gene genealogy. BMC Evolutionary Biology 8, 1.

66. Delmas, S., Shunburne, L., Ngo, H.P., and Allers, T. (2009). Mre11-Rad50 promotes rapid repair of DNA damage in the polyploid archaeon Haloferax volcanii by restraining homologous recombination. PLoS Genet 5, e1000552.

67. Rogers, R.L., Bedford, T., and Hartl, D.L. (2009). Formation and longevity of chimeric and duplicate genes in Drosophila melanogaster. Genetics 181, 313–322.

68. Yang, S., Arguello, J.R., Li, X., Ding, Y., Zhou, Q., Chen, Y., Zhang, Y., Zhao, R., Brunet, F., Peng, L., et al. (2008). Repetitive element-mediated recombination as a mechanism for new gene origination in Drosophila. PLoS Genet 4, e3.

69. Kashkush, K., Feldman, M., and Levy, A.A. (2003). Transcriptional activation of retrotransposons alters the expression of adjacent genes in wheat. Nature genetics 33, 102–106.

70. Kenan-Eichler, M., Leshkowitz, D., Tal, L., Noor, E., Melamed-Bessudo, C., Feldman, M., and Levy, A.A. (2011). Wheat hybridization and polyploidization results in deregulation of small RNAs. Genetics 188, 263–272.

71. Levin, H.L., and Moran, J.V. (2011). Dynamic interactions between transposable elements and their hosts. Nature Reviews Genetics 12, 615–627.

72. Liu, D., You, C., Liu, S., Liu, L., Duan, W., Chen, S., Yan, J., and Liu, Y. (2009). Characterization of a novel Tc1-like transposon from bream (Cyprinidae, Megalobrama) and its genetic variation in the polyploidy progeny of bream-red crucian carp crosses. Journal of molecular evolution 69, 395–403.

73. Shapiro, J.A. (2005). A 21st century view of evolution: genome system architecture, repetitive DNA, and natural genetic engineering. Gene 345, 91–100.

74. Collins, S., and Bell, G. (2004). Phenotypic consequences of 1,000 generations of selection at elevated CO2 in a green alga. Nature 431, 566–569.

75. Edelist, C., Raffoux, X., Falque, M., Dillmann, C., Sicard, D., Rieseberg, L.H., and Karrenberg, S. (2009). Differential expression of candidate salt-tolerance genes in the halophyte Helianthus paradoxus and its glycophyte progenitors H. annuus and H. petiolaris (Asteraceae). American Journal of Botany 96, 1830–1838.

76. Groszmann, M., Greaves, I.K., Albertyn, Z.I., Scofield, G.N., Peacock, W.J., and Dennis, E.S. (2011). Changes in 24-nt siRNA levels in Arabidopsis hybrids suggest an epigenetic contribution to hybrid vigor. Proceedings of the National Academy of Sciences of the United States of America 108, 2617–2622.

77. Arnold, M., Ballerini, E., and Brothers, A. (2012). Hybrid fitness, adaptation and evolutionary diversification: lessons learned from Louisiana Irises. Heredity 108, 159–166.

78. Tirosh, I., Reikhav, S., Levy, A.A., and Barkai, N. (2009). A Yeast Hybrid Provides Insight into the Evolution of Gene Expression Regulation. Science 324, 659–662.

79. Rieseberg, L.H. (2001). Chromosomal rearrangements and speciation. Science 16, 351–358.

80. Nei, M., and Nozawa, M. (2011). Roles of mutation and selection in speciation: from Hugo de Vries to the modern genomic era. Genome biology and evolution 3, 812–829.

81. Koyama, H., Ito, T., Nakanishi, T., and Sekimizu, K. (2007). Stimulation of RNA polymerase II transcript cleavage activity contributes to maintain transcriptional fidelity in yeast. Genes to Cells 12, 547–559.

82. Perathoner, S., Daane, J.M., Henrion, U., Seebohm, G., Higdon, C.W., Johnson, S.L., Nusslein-Volhard, C., and Harris, M.P. (2014). Bioelectric signaling regulates size in zebrafish fins. PLoS Genet 10, e1004080.

83. Xu, K., Wen, M., Duan, W., Ren, L., Hu, F., Xiao, J., Wang, J., Tao, M., Zhang, C., and Wang, J. (2015). Comparative Analysis of Testis Transcriptomes from Triploid and Fertile Diploid Cyprinid Fish. Biology of reproduction, biolreprod. 114. 125609.

84. Zhou, Y., Zhong, H., Liu, S., Yu, F., Hu, J., Zhang, C., Tao, M., and Liu, Y. (2014). Elevated expression of Piwi and piRNAs in ovaries of triploid crucian carp. Molecular and cellular endocrinology 383, 1–9.

